# Ribosome profiling elucidates differential gene expression in bundle sheath and mesophyll cells in maize

**DOI:** 10.1101/2020.12.15.422948

**Authors:** Prakitchai Chotewutmontri, Alice Barkan

**Affiliations:** Institute of Molecular Biology, University of Oregon, Eugene, OR 97403 USA

## Abstract

The efficiencies offered by C_4_ photosynthesis have motivated efforts to understand its biochemical, genetic and developmental basis. Reactions underlying C_4_ traits in most C_4_ plants are partitioned between two cell types, bundle sheath (BS) and mesophyll (M) cells. RNA-seq has been used to catalog differential gene expression in BS and M cells in maize and several other C_4_ species. However, the contribution of translational control to maintaining the distinct proteomes of BS and M cells has not been addressed. In this study, we used ribosome profiling (ribo-seq) and RNA-seq to describe translatomes, translational efficiencies, and microRNA abundance in BS and M-enriched fractions of maize seedling leaves. A conservative interpretation of our data revealed 182 genes exhibiting cell-type dependent differences in translational efficiency, 31 of which encode proteins with core roles in C_4_ photosynthesis. Our results suggest that non-AUG start codons are used preferentially in upstream open reading frames of BS cells, revealed mRNA sequence motifs that correlate with cell type-dependent translation, and identified potential translational regulators that are differentially expressed. In addition, our data expand the set of genes known to be differentially expressed in BS and M cells, including genes encoding transcription factors and microRNAs. These data add to the resources for understanding the evolutionary and developmental basis of C_4_ photosynthesis and for its engineering into C_3_ crops.

## Introduction

Plant species are classified as C_3_ or C_4_ according to their mechanism of photosynthetic carbon fixation. C_4_ photosynthesis offers advantages under hot, dry conditions (Hatch, 1987), and it evolved many times from a C_3_ progenitor (Sage et al., 2011; Schlüter and Weber, 2020). Many C_4_ plants are characterized by a specialized leaf anatomy, denoted Kranz anatomy, that partitions enzymes of the C_4_ pathway between two cell types: mesophyll (M) and bundle sheath (BS). M cells surround BS cells, which in turn surround vascular bundles. Atmospheric CO_2_ is first fixed by phosphoenolpyruvate (PEP) carboxylase in M cells to produce four-carbon acids. These are transported to BS cells, where they are decarboxylated, providing CO_2_ for fixation by ribulose bisphosphate carboxylase/oxygenase (Rubisco) according to the C_3_ scheme. This organization reduces Rubisco’s wasteful oxygenation reaction by increasing local CO_2_ concentration and decreasing local O_2_ concentration.

Research into the genetic and developmental mechanisms underlying C_4_ traits has been propelled by interest in introducing C_4_ traits into C_3_ crops (Sedelnikova et al., 2018; Ermakova et al., 2020). A thorough understanding of patterns of gene expression during C_4_ differentiation is essential to achieve this goal. Numerous studies have approached this problem through transcriptome analysis (Wang et al., 2016; Schlüter and Weber, 2020), including studies that profiled transcriptomes during development of C_4_ leaves (Li et al., 2010; Pick et al., 2011; Liu et al., 2013; Wang et al., 2013b; Wang et al., 2014; Ding et al., 2015; Mattiello et al., 2015; Denton et al., 2017), or compared transcriptomes between BS and M cells (Li et al., 2010; Chang et al., 2012; Aubry et al., 2014; John et al., 2014; Tausta et al., 2014; Denton et al., 2017), among different C_4_ lineages (Aubry et al., 2014; Chen et al., 2014; John et al., 2014; Ding et al., 2015; Li et al., 2015; Offermann et al., 2015; Covshoff et al., 2016; Rao et al., 2016) or among related C_3_ and C_4_ species (Bräutigam et al., 2011; Gowik et al., 2011; Chen et al., 2014; Külahoglu et al., 2014; Wang et al., 2014; Ding et al., 2015; Lauterbach et al., 2017; Schlüter et al., 2017; Dunning et al., 2019). These transcriptome data have been complemented by surveys of protein populations during C_4_ differentiation and in isolated BS and M chloroplasts (Majeran et al., 2005; Bräutigam et al., 2008; Majeran et al., 2008; Friso et al., 2010; Majeran et al., 2010).

The results from these studies support the view that transcriptional control plays the major role in determining patterns of gene expression during C_4_ differentiation and in mature BS and M cells. That said, evidence for post-transcriptional contributions emerged in several studies (John et al., 2014; Ponnala et al., 2014; Williams et al., 2016), yet translational regulation has barely been explored (Schlüter and Weber, 2020). Ribosome profiling (ribo-seq) provides the means to comprehensively address this issue. Ribo-seq uses deep sequencing to map and quantify ribosome-protected mRNA fragments (ribosome footprints). Because average translation elongation rates are generally similar among mRNAs under a given condition, the normalized abundance of ribosome footprints is a widely accepted proxy for relative rates of protein synthesis (Brar and Weissman, 2015). Comparison of ribosome footprint abundance with the abundance of the corresponding mRNA allows inferences about translational efficiencies on a genome-wide scale.

Previously, we used ribo-seq in conjunction with RNA-seq to analyze chloroplast gene expression in BS-and M-enriched leaf fractions in maize, a C_4_ species of the NADP-malic enzyme (NADP-ME) subtype (Chotewutmontri and Barkan, 2016). We found that differences in mRNA abundance largely account for differential expression of chloroplast genes in the two cell types, but differences in translational efficiency synergize with mRNA-level effects in some cases. We have now extended this analysis to cytosolic mRNAs. Similar to what we observed in chloroplasts, the differential expression of nuclear genes in BS and M cells results primarily from differences in mRNA abundance. However, our data identified a subset of mRNAs whose translational efficiencies are significantly different in the two cell types, and revealed mRNA sequence features and *trans-*factors that correlate with cell-type dependent translation. Additionally, our results expand the set of genes known to be differentially expressed in maize BS and M cells, suggesting new genes of potential relevance to C_4_ traits.

## Results

### Overview of ribo-seq and RNA-seq data collected from BS- and M-enriched leaf fractions

Analyses reported here used BS- and M-enriched fractions generated from the apical region of seedling leaves with a rapid mechanical fractionation method similar to that used in our prior study of chloroplast translatomes (Chotewutmontri and Barkan, 2016). Marker proteins for each cell type were highly enriched in these fractions (Fig. 1A). Ribosome footprints and RNA were purified from aliquots of the same BS and M preparations, in three biological replicates. The ribo-seq reads mapping to cytosolic mRNAs exhibit the expected characteristics of ribosome footprints: they map almost exclusively to protein coding regions, their size distribution is heavily weighted to 29-30 nucleotides, and their positions within open reading frames (ORFs) show strong 3-nucleotide periodicity (Supplemental Fig. S1A-C). We used rRNA-depleted total RNA for RNA-seq to avoid the 3’-bias that can lead to false inferences about differential expression (Denton et al., 2017) (Supplemental Fig. S2A-B). Correlation coefficients among replicates ranged from approximately 0.93 to 0.98, with replicate datasets clustering together as expected (Supplemental Fig. S1D).

**Figure 1.**
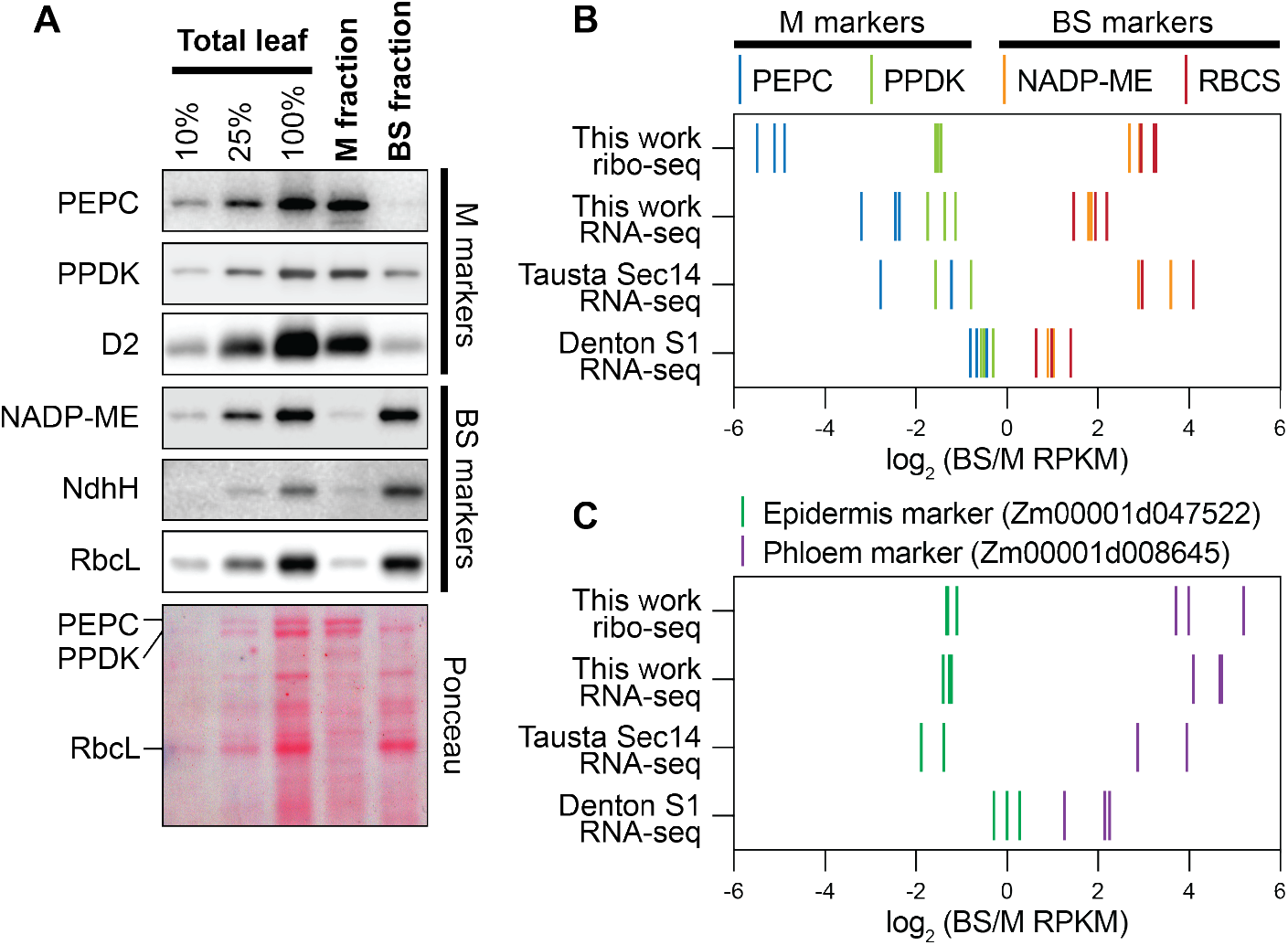
Enrichment of cell-type specific markers in BS and M fractions. (A) Immunoblots showing abundance of marker proteins in BS and M fractions. Replicate blots were probed with the indicated antibodies. BS markers: NADP-dependent malic enzyme (NADP-ME), H subunit of the NDH-like complex (NdhH), Rubisco large subunit (RbcL). M markers: phosphoenolpyruvate carboxylase (PEPC), pyruvate orthophosphate dikinase (PPDK), D2 subunit of Photosystem II (D2). An image of one blot stained with Ponceau S is shown below. (B) Abundance of RNA-seq and ribo-seq reads from BS and M marker genes. BS markers: NADP-ME (Zm00001d000316) and Rubisco small subunit (RBCS; Zm00001d052595). M markers: PEPC (Zm00001d046170) and PPDK (Zm00001d038163). RNA-seq data from Tausta et al (2014) (leaf section 14 with 2 replicates) and Denton et al (2017) (leaf section S1 with 3 replicates) are compared with our data. Each line symbol represents normalized reads for one gene (see color key at top) in one replicate. RPKM, reads per kilobase per million reads mapped to nuclear coding sequences. (C) Abundance of RNA-seq and ribo-seq reads from epidermis and phloem marker genes. Data for additional marker genes are shown in Supplemental Fig. S2C.

Four RNA-seq studies of maize BS and M fractions were reported previously (Li et al., 2010; Chang et al., 2012; Tausta et al., 2014; Denton et al., 2017), all of which used polyA-selected mRNA. We used the two most recent studies as points of comparison for our data. Tausta et al (Tausta et al., 2014) generated BS and M-enriched fractions by laser capture microdissection whereas Denton et al (Denton et al., 2017) used a mechanical method that differed from ours. Both studies analyzed tissue slices at several positions along the leaf blade, which represent different stages along the pathway of photosynthetic differentiation. The apical sections were most analogous to our material, so we selected those data for our comparisons. To facilitate comparisons, we aligned both prior datasets to the current maize genome assembly (B73 RefGen_v4) and we calculated differential expression using the same analysis pipeline we used for our data.

Figure 1B compares the cell type-enrichment of sequence reads from four C_4_ marker genes in our data to those in the prior datasets. In comparison with the Denton data, our RNA-seq data exhibited greater enrichment of both M and BS markers. In comparison with the Tausta data, our RNA-seq data exhibited similar enrichment of M markers and less enrichment of BS markers. We expected that our mechanical fractionation method would result in contamination of the M and BS fractions with epidermal and vascular tissue, respectively, and analysis of epidermal and phloem markers confirmed this to be the case (Fig. 1C and Supplemental Fig. S2C). Despite the considerable differences in sample preparation and developmental stage, a core gene set associated with C_4_ photosynthesis (Schlüter and Weber, 2020) showed Pearson correlation coefficients of at least 0.85 among all three datasets (Supplemental Fig. S3A).

Comparison of our ribo-seq and RNA-seq data (Supplemental Fig. S3B) returned correlation coefficients of 0.72 and 0.89 for genome-wide and C_4_ gene comparisons, respectively. These values are considerably lower than those for replicate samples, suggesting some differences in translational efficiency in the two cell types. In addition, we compared our data to that from a proteomic study of material obtained with a similar mechanical fractionation method (Friso et al., 2010) (Supplemental Fig. S3C). Proteins that were quantified with higher confidence levels showed considerable correlation with our ribo-seq data. Furthermore, our ribo-seq data correlated better with the proteomic data than did our RNA-seq data, consistent with a role for translational control in maintaining the distinct proteomes in the two cell types.

### Differences in translational efficiency contribute to the differential expression of genes in BS and M cells

To detect mRNAs that experience differential translation in M and BS cells, we calculated differences in translational efficiency (TE) with XTAIL (Xiao et al., 2016). XTAIL reports the normalized ratio of ribo-seq to RNA-seq reads for each gene together with a false discovery rate (FDR) for differential translation. We drew conclusions about differential translation only for genes with an average of at least 100 RNA-seq reads mapped to coding sequences in both the BS and M fractions, as well as an average of at least 100 ribo-seq reads in either the BS or M fraction. 9,476 genes met these read-count criteria. The XTAIL output for these genes is provided in Supplemental Data S1. We defined a gene as being differentially translated using stringent criteria: > 3-fold difference in TE between the two tissues and an FDR < 0.001. 355 genes met these criteria (Fig. 2A, Supplemental Data S2). A comparison of the ribo-seq and RNA-seq data (Fig. 2B) parses these genes into several groups: those that are differentially expressed primarily due to differences in translational efficiency (TE), those for which changes in RNA and TE synergize to amplify differential expression (RNA&TE), and those for which a change in TE offsets a change in RNA (buffering). Buffering of this type has been detected in many ribo-seq studies (McManus et al., 2014; Schafer et al., 2015; Oertlin et al., 2019). However, we are hesitant to draw conclusions about the buffering set because our RNA-seq data for those particular genes correlated poorly with data from prior RNA-seq studies (Supplemental Fig. S4). By contrast, RNA-seq data for our TE and RNA&TE sets correlated well with those from prior studies (Supplemental Fig. S4). Examples of read coverage plots for genes whose cell-type specific expression results primarily from a difference in TE are shown in Figure 2C with additional examples in Supplemental Figure S5.

**Figure 2.**
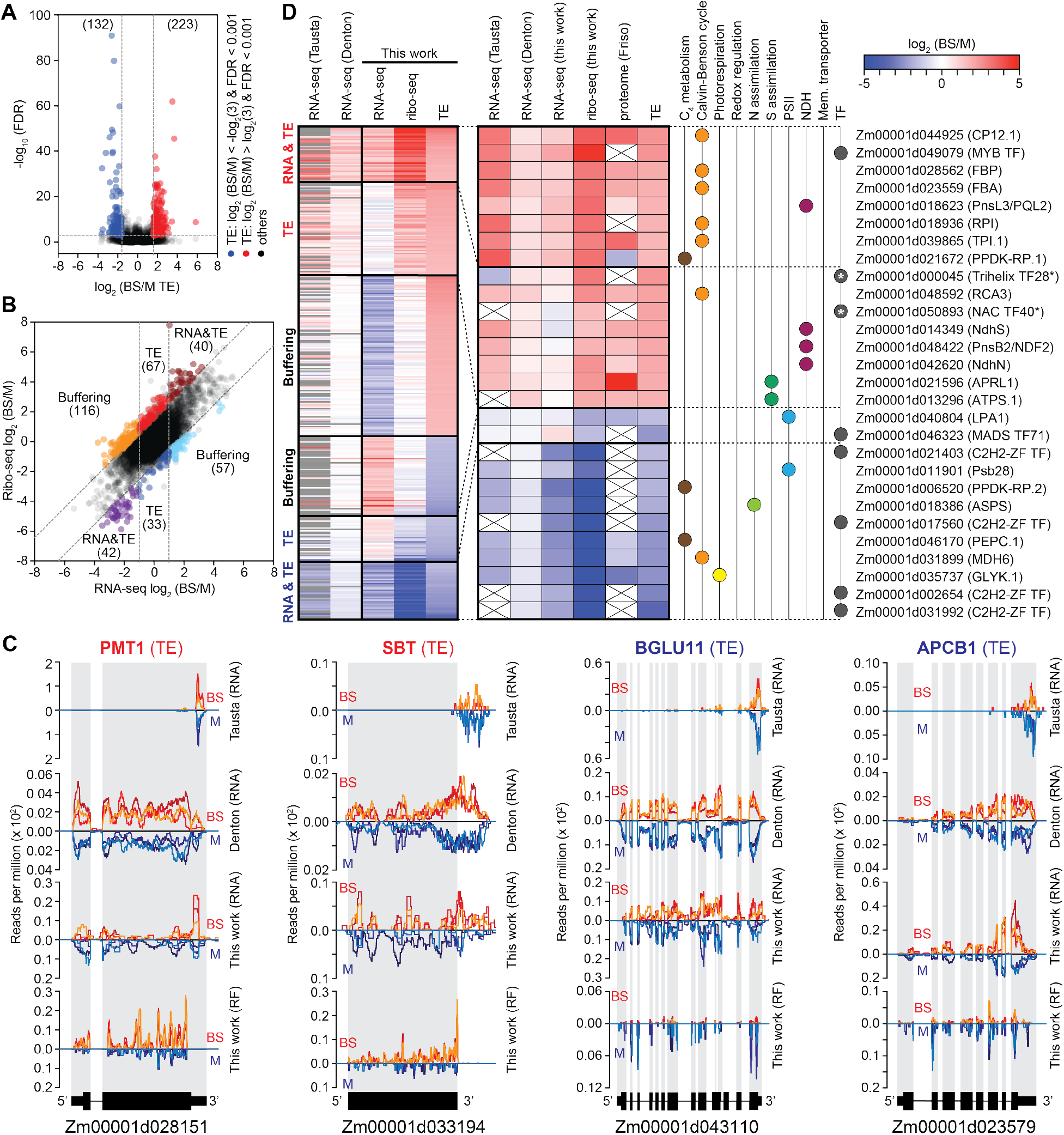
Overview of genes exhibiting differential translation in BS and M fractions. (A) Volcano plot of translational efficiency (TE) and FDR values. Each symbol represents one gene. Genes exhibiting preferential translation in M or BS fractions are shown in blue or red, respectively. Dashed lines indicate the FDR and fold-change cutoffs we used to define differentially translated genes. (B) Scatter plot comparing RNA-seq to ribo-seq BS/M ratios. Each symbol represents one gene. Gene sets are labeled according to the dominant feature underlying differential expression (TE, RNA, or both). Dashed lines indicate the fold-change cutoffs we used to categorize the genes. (C) Examples of RNA-seq and ribosome footprint (RF) read coverage for genes whose differential expression results primarily from differential translation. Normalized read coverage in BS or M fractions is shown above or below the line, respectively. Coverage was normalized by million reads mapped to nuclear coding sequences. Each line represents one replicate. The re-analyzed values of published RNA-seq data are shown for comparison. Additional examples are shown in Supplemental Figure S5. (D) Heatmaps showing BS/M values for genes exhibiting differential translation. Prior RNA-seq [Tausta et al (2014) leaf section 14 and Denton et al (2017) leaf section S1] and proteome (Friso et al (2010) data are shown for comparison. The previous RNA-seq data were re-aligned with our pipeline and analyzed with DESeq2; genes that did not meet our read count cutoffs are in gray. The expanded heat map to the right provides information for genes that are of particular relevance to C4 photosynthesis. Asterisks mark transcription factors (TF) for which differential TE plays the dominant role in dictating cell type-specific expression that were not reported as differentially expressed in RNA-seq studies. PSII, Photosystem II; NDH, NADH dehydrogenase-like complex; Mem., membrane. Values are provided in Supplemental Data S2.

The data for translationally regulated genes of particular relevance to C_4_ physiology are shown in an expansion of the heat map in Figure 2D. These include (i) “C_4_” genes (Schlüter and Weber, 2020) involved in photorespiration, the Calvin-Benson cycle, C_4_ metabolism, redox regulation, and nitrogen and sulfur assimilation; (ii) membrane transporters, subunits of PSII, and subunits of the NADH dehydrogenase-like complex (NDH) that are strongly enriched in one cell type or the other (Majeran and van Wijk, 2009); and (iii) genes encoding transcription factors, which could potentially contribute to the establishment of cell-type specific transcriptomes. Roughly one third of these genes are regulated primarily at the level of translational efficiency (TE), and the remainder by a combination of RNA abundance and TE. The latter set includes the gene encoding ribose-5-phosphate isomerase (RPI), which had been suggested to be under translational control based on a comparison of Setaria and maize transcriptome data (John et al., 2014). We detected three transcription factors that are differentially expressed primarily due to a difference in translational efficiency, two of which (Trihelix TF28 and NAC TF40) had not been detected as differentially expressed in transcriptome studies (Fig. 2D asterisks).

### Features of untranslated regions that correlate with differential translation in BS and M cells

The rate of translation initiation is influenced by various features of mRNA untranslated regions (UTRs), including upstream open reading frames (uORFs), start codon sequence and sequence context, RNA structure, and binding sites of translational regulators. To gain insight into the basis for differences in translational efficiency between BS and M cells, we compared UTR features among mRNAs exhibiting cell-type dependent translation to those that do not. RNAs that are translated preferentially in the BS fraction show a tendency toward UTRs that are shorter and more GC-rich than those in the control set, particularly in the 5’ UTR (Fig. 3A). We identified six sequence motifs that are enriched in UTRs of mRNAs that are preferentially translated in one cell type or the other (Fig. 3B and Supplemental Data S3): three in 5’ UTRs of the BS set, one in 3’ UTRs of the BS set, and one each in 5’ and 3’ UTRs of the M set. Out of the 182 genes whose cell-type dependent translation contributes to differential expression (>3-fold difference in TE), 78 harbor at least one of these motifs and 20 harbor more than one (Fig. 3C). Enriched sequence motifs of this nature are candidates for binding sites of translational regulators. Few sequence-specific RNA binding proteins have been identified in plants that regulate the translation of specific mRNAs. However, members of the PUF (Pumilio/FBF) family are attractive candidates because they are known to regulate translation and RNA stability by binding specific UTRs in animals and fungi (Goldstrohm et al., 2018; Wang et al., 2018), and the PUF family is unusually large in plants (Dedow and Bailey-Serres, 2019; Joshna et al., 2020). Furthermore, the UGUGCCG motif that is enriched in 3’ UTRs of the BS set (Fig. 3B) resembles the consensus UGUR core binding site for canonical PUF proteins. With this in mind, we examined expression of genes encoding PUF proteins in BS and M fractions in our ribo-seq and RNA-seq data and in prior RNA-seq studies (Fig. 3D). We identified several PUF-encoding genes that are expressed preferentially in one cell type or the other in at least one dataset (Fig. 3D). A particularly intriguing example is Zm00001d023781, whose expression is highly enriched in BS cells across multiple datasets. Furthermore, the expression of this gene along the seedling leaf blade peaks in the zone of active chloroplast biogenesis (http://bar.utoronto.ca/efp_maize/cgi-bin/efpWeb.cgi) (Li et al., 2010; Wang et al., 2014), consistent with a role in regulating chloroplast differentiation. These results highlight Zm00001d023781 as an attractive candidate for future investigation with regard to the post-transcriptional control of BS and M differentiation.

**Figure 3.**
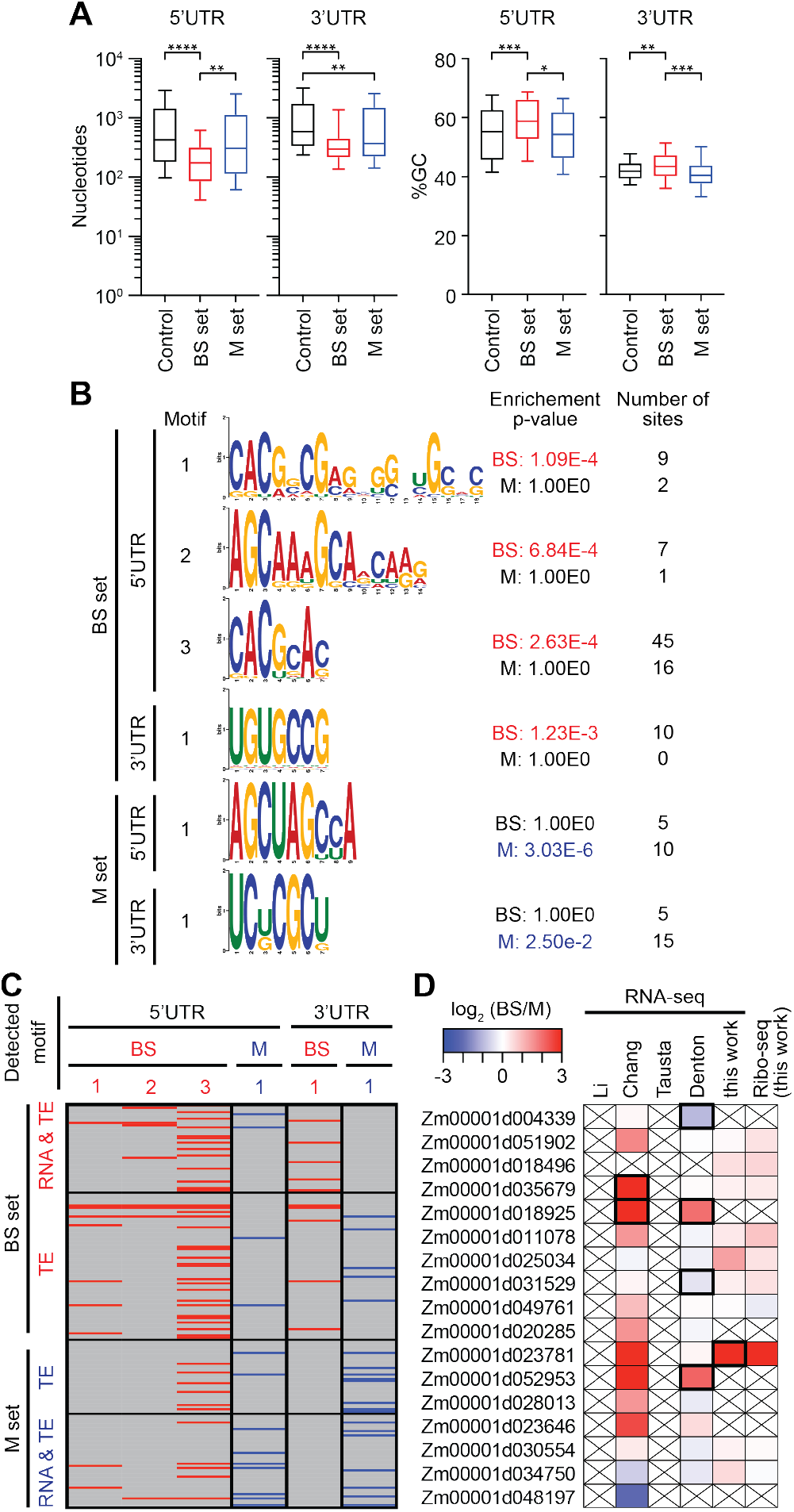
Characteristics of untranslated regions of differentially translated genes. Differentially translated genes used for these analyses are the TE and RNA&TE sets from Figure 2B (|log_2_ (BS/M TE)| > 1.585, FDR < 0.001 excluding buffering sets, BS set n=107 and M set n=75). The control set was defined as |log_2_ (BS/M) TE| ≤ 0.585 (n=5280). (A) UTR length and GC content. Horizontal lines show median values, boxes show 25^th^ to 75^th^ percentile, and whiskers indicate 10th to 90th percentiles. Brackets show significant p-values from Dunn’s Multiple Comparison Test. * p=0.01; ** p<0.005; *** p=0.002, **** p<0.0001. (B) Enriched motifs in UTRs of differentially translated genes. Enriched motifs were identified with STREME (Bailey, 2020) and those matching known DNA motifs were removed. Motifs reported by AME (McLeay and Bailey, 2010) as significantly enriched (p-value < 0.05) are shown. (C) Presence of enriched motifs in differentially translated genes. Gene sets are taken from Figure 2B. Motifs are identified at top via the motif numbers shown in panel (B). (D) Differential expression of PUF genes. Values from prior RNA-seq datasets come from Li et al (2010), Chang et al (2012), Tausta et al (2014) section 14 and Denton et al (2017) section S1. Genes reported with high confidence as differentially expressed in prior studies or in our data [|log_2_ (BS/M)| > 1, FDR < 0.001] are bordered in black. Crossed boxes indicate absence of data or read counts below cutoffs.

Upstream open reading frames (uORFs) often regulate the translation of main ORFs (mORFs) in eukaryotic mRNAs (Hinnebusch et al., 2016). We inspected the 335 genes for which TE differed by at least 2-fold in the BS and M fractions (excluding the buffering set) for ribosome footprints upstream of their annotated start codon. We considered uORFs beginning with AUG as well as those with the non-canonical start codons CUG and ACG (van der Horst et al., 2019). Thirty-nine genes fit these criteria (Supplemental Data S4): five had only uORFs beginning with AUG, 28 had uORFs beginning with CUG or ACG, six had both AUG and non-AUG uORFs, nine had overlapping uORFs, and 17 had uORFs that overlap mORFs.

Comparison of uORF and mORF ribosome occupancies (ribo-seq/RNA-seq) in BS and M fractions revealed three types of relationship (Fig. 4A and Supplemental Data S4): uORF translation was inversely related to mORF translation, suggesting a repressive effect of the uORF; uORF translation was positively correlated with mORF translation, suggesting an activating effect of the uORF; and uORF and mORF translation changed independently of one another. Figure 4B shows examples of normalized ribo-seq read coverage from two genes in each category. Interestingly, our uORF data suggest a bias toward translation of non-AUG start codons in BS cells (Fig. 4C). However, a deeper analysis of start codon usage will be needed to draw firm conclusions. In yeast, the translation initiation factors eIF1 and eIF5A promote translation from AUG codons while eIF5 antagonizes this function, thereby favoring non-AUG initiation (Hinnebusch et al., 2016; Eisenberg et al., 2020). We therefore considered the possibility that differing ratios of these factors in BS and M cells underlie the differences in AUG versus non-AUG uORF initiation suggested by our data. In fact, the ratio of eIF1/eIF5A to eIF5 expression is elevated in BS cells (Fig. 4D), running counter to the hypothesized relationship with non-AUG initiation. In particular, expression of one eIF1 paralog (Zm00001d021668) and two eIF5A paralogs (Zm00001d006760, Zm00001d022042) is BS-enriched in our ribo-seq data (Fig. 4D) and in many prior RNA-seq datasets (Supplemental Fig. S6), suggesting that these paralogs may play non-canonical roles that favor non-AUG initiations.

**Figure 4.**
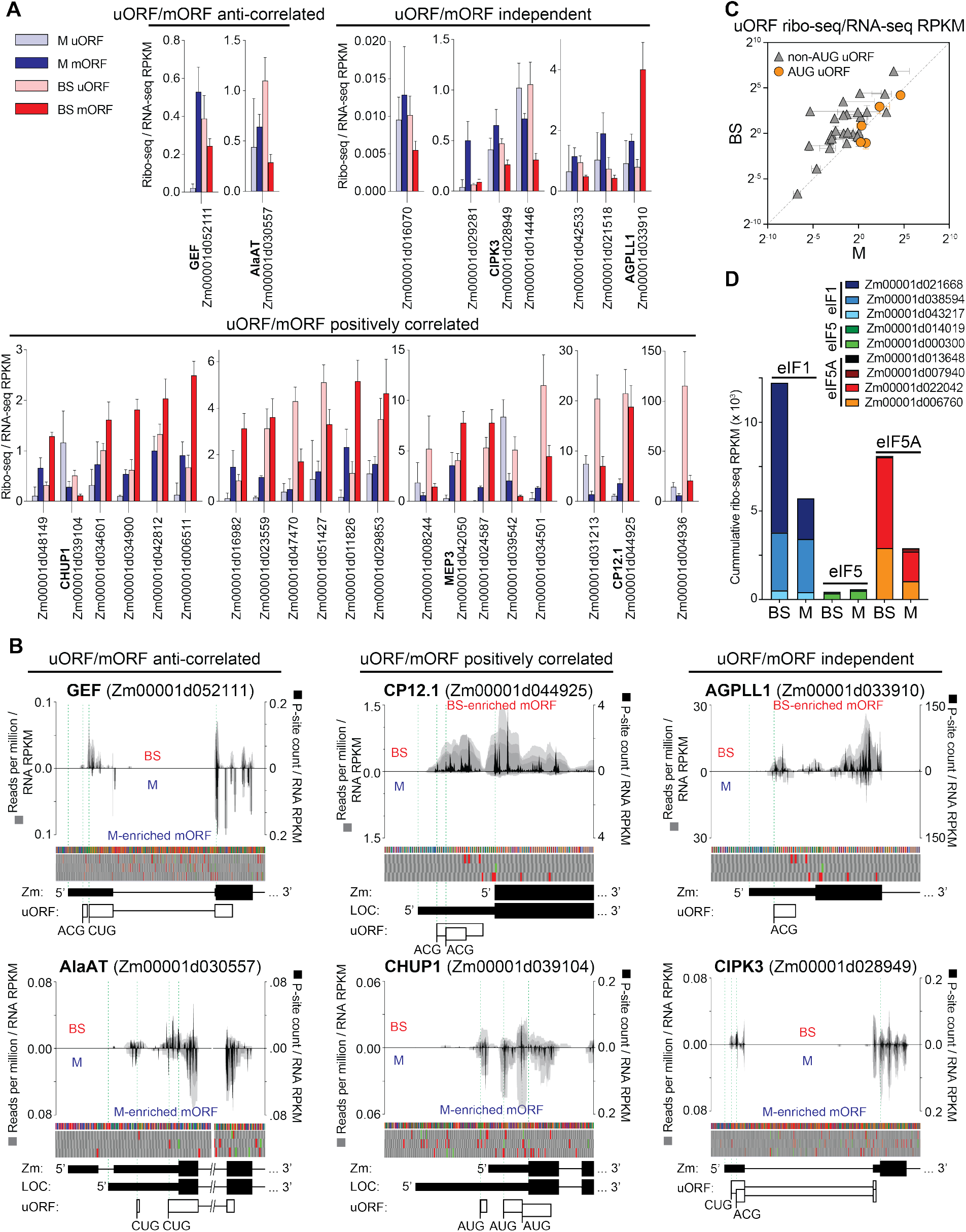
Translation of uORFs of differentially translated genes. The genes used for this analysis had |log_2_ (BS/M) TE| > 1 and FDR < 0.001, excluding the buffering sets. Ribo-seq read counts exclude ribosomes bound to the mORF start codon. RPKM, reads per kilobase per million reads mapped to nuclear coding sequences. (A) TE (ribo-seq/RNA-seq RPKM) of uORFs and mORFs in BS and M fractions. Genes are grouped based on whether differences in TE of the uORF and mORF in BS and M fractions are correlated, anti-correlated, or independent. Thirty-nine differentially translated genes had ribosomes in uORFs; data are shown for the 29 genes with uORFs that begin with either AUG or non-AUG codons (but not both) and for which the relationship between uORF and mORF translation was reproducible. Data for the other ten genes are summarized in Supplemental Data S4. (B) Examples of normalized ribo-seq read coverage from the 5’ regions of differentially translated uORF-containing genes. Gray shading indicates coverage from each replicate such that overlapping coverage from different replicates appears as a darker shade. Coverage in BS or M fractions is shown above or below the line, respectively. Values were normalized by million ribo-seq reads mapped to nuclear coding sequence and by RNA-seq RPKM for the mORF. Normalized P-site occupancies (sum of replicates) are shown in black (scale to the right). Nucleotide and 3-frame translation tracks are shown above gene models. The green and red boxes in the translation tracks indicate AUG start and stop codons, respectively. Zm and LOC indicate gene models from Gramene and GenBank, respectively. (C) TE (ribo-seq/RNA-seq) of uORFs with AUG or non-AUG start codons in BS and M fractions. Each symbol represents one gene. Values are the mean ± SD from three replicates. (D) Expression of eIF1, eIF5 and eIF5A in BS and M fractions based on ribo-seq data (sum of three replicates).

### Global analysis of BS and M translatomes expands the known set of differentially expressed genes

The abundance of ribosome footprints mapping to a gene reflects both mRNA abundance and translational efficiency. This is widely used as a proxy for relative rates of protein synthesis among genes, and there is considerable evidence that this is, with rare exceptions, a valid assumption (Brar and Weissman, 2015). Therefore, ribo-seq can provide a more accurate picture of differential expression than RNA-seq. With that in mind, we analyzed our ribo-seq data with DESeq2 to catalog genes that are differentially expressed in BS and M cells. We limited the analysis to the 13,182 genes having an average of at least 100 ribo-seq reads mapped to coding sequences in at least one of the two tissue types (Supplemental Data S5). We defined differentially expressed genes as having |log_2_ (BS/M)| > 1 and FDR < 0.001. With these criteria, we detected 1,345 and 993 genes that are preferentially expressed in the BS and M fractions, respectively (Fig. 5A, Supplemental Data S6 and S7). We then compared these results to those from previous RNA-seq studies of maize BS and M fractions (Li et al., 2010; Chang et al., 2012; Tausta et al., 2014; Denton et al., 2017). Approximately 30% of the genes reported as differentially expressed in our ribo-seq analysis were not reported to be differentially expressed in prior RNA-seq studies (Supplemental Data S6 and S7; “unique” sets in Fig. 5A). These “unique” gene sets showed similar functional enrichment patterns as the complete sets of BS-and M-enriched genes (Fig. 5B, Supplemental Data S8), instilling confidence that many of these genes are, in fact, differentially expressed in BS and M cells. The unique set includes many transcription factors (Fig. 5C), as discussed below.

**Figure 5.**
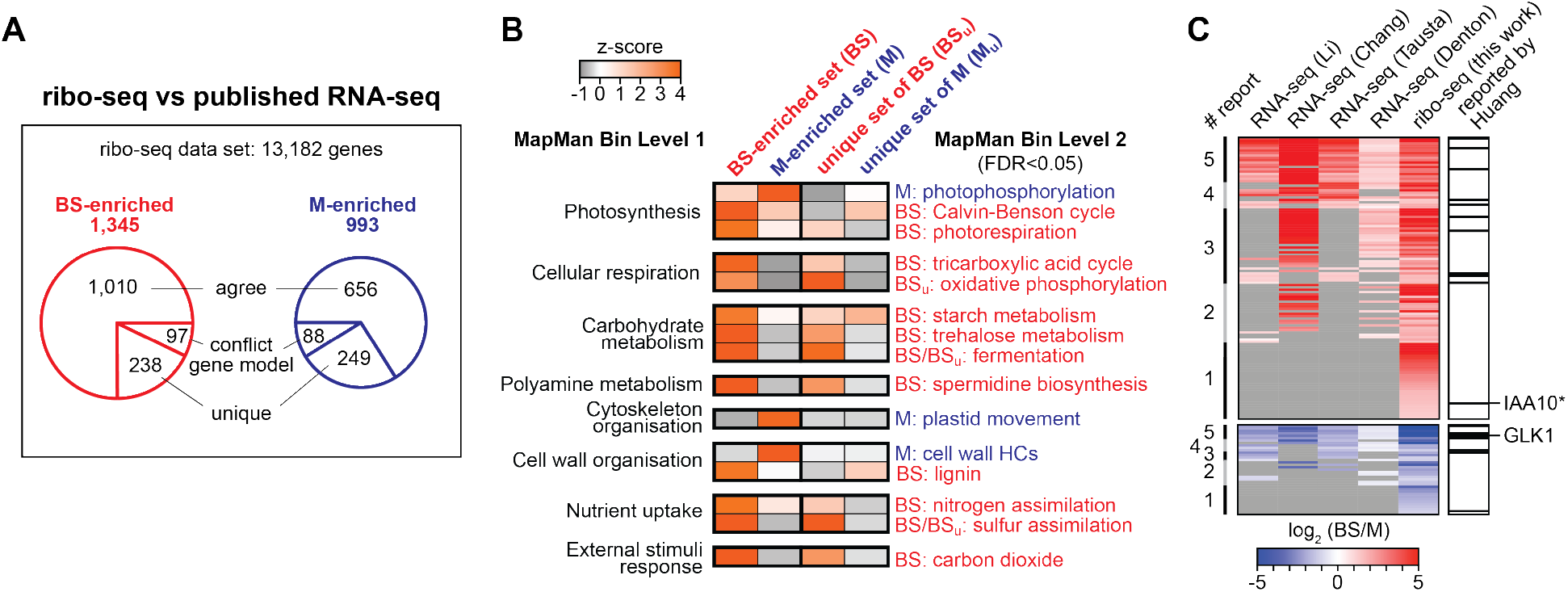
Comparison of translatomes in BS and M fractions. (A) Overview of ribo-seq data. 13,182 genes met the read-count cutoff used in this analysis (average of ≥100 reads in at least one of the two fractions). The BS-and M-enriched gene sets [|log_2_ (BS/M)| >1 and FDR < 0.001] were compared to genes that had been reported to be differentially expressed in previous RNA-seq studies: Chang et al (2012), Li et al (2010), Denton et al (2017) section S1, Tausta et al (2014) section 14. The “conflict gene model” sets consist of genes that cannot be compared among studies because of conflicting gene models. The “agree” sets include genes that were reported by at least one prior RNA-seq study to be preferentially expressed in the same tissue type as in our ribo-seq data. The “unique” sets are genes reported by our ribo-seq data as differentially expressed that were not reported as differentially expressed in prior studies. (B) Functional enrichment analysis of genes identified as differentially expressed in our ribo-seq data that had not been reported as differentially expressed in previous studies. Heatmaps display the term enrichment z-scores when comparing BS-and M-enriched gene sets to the set of all expressed genes. Data for the complete BS and M-enriched sets (BS and M) are compared with data for the unique sets described in (A). HC, hydroxycinnamic acid. (C) Heatmap of log_2_ (BS/M) values of ribo-seq data for transcription factor (TF) genes that exhibited BS or M-enriched ribosome footprint abundance. Values from prior RNA-seq studies are shown for comparison. Genes are ranked based on the number of reports concluding that the gene is differentially expressed (# report, left). Genes not reported as differentially expressed are in gray. The TFs that were identified in a cross-species transcriptome analysis by Huang et al (2016) are marked in the right column. Transcription factors reported to be differentially expressed among all studies are listed in Supplemental Data S10. IAA10*, Aux/IAA-transcription factor 10 also known as ZmIAA7; GLK1, Golden2-like 1.

Three photosynthetic complexes harboring plastid-encoded subunits accumulate differentially in BS and M cells: Rubisco and the NADH dehydrogenase-like complex (NDH) accumulate preferentially in BS cells whereas PSII accumulates preferentially in M cells. The biogenesis of each complex involves a large number of nuclear genes, some of which are involved in complex assembly and others in the expression of plastid-encoded subunits. Differential expression data for genes that function specifically in the biogenesis of these complexes (Supplemental Fig. S7 and Supplemental Data S9) show that many of them are preferentially expressed in the same cell type as the cognate complex. Furthermore, three genes encoding proteins that activate translation of the chloroplast *psbA* mRNA (HCF173, HCF244, OHP2) (Schult et al., 2007; Link et al., 2012; Chotewutmontri et al., 2020) are expressed preferentially in M cells. This can account for the preferential translation of *psbA* mRNA in M cells we had reported previously (Chotewutmontri and Barkan, 2016).

### Differential expression of transcription factors and microRNAs in BS and M fractions

The distinct transcriptomes of BS and M cells are presumably maintained, at least in part, by cell-type specific regulatory molecules, including transcription factors and microRNAs. We identified genes encoding 114 and 37 transcription factors that were enriched in BS and M translatomes, respectively (Supplemental Data S10). Approximately one third of these had not been reported as differentially expressed in prior RNA-seq studies. One interesting example is a BS-enriched member of the Aux/IAA family of auxin-regulated transcription factors (Gene ID Zm00001d041416; IAA10 in Fig. 5C), which also emerged as relevant to C_4_ biology based on evolutionary inferences (Huang et al., 2017). Aux/IAA proteins govern diverse developmental processes including vascular patterning (Zhang et al., 2014; Luo et al., 2018). These observations suggest the possibility that this particular Aux/IAA protein regulates C_4_ vascular patterning in the maize leaf. Only one fifth of these transcription factors were reported as differentially expressed across all studies (Fig. 5C). The low concordance among studies is likely due to a variety of factors, including varying sequencing depth, varying criteria used to identify differentially expressed genes, and imperfectly matched developmental stages. In addition, cell-type dependent translation accounted for the differential expression of two of the transcription factors detected uniquely in our set (Trihelix TF28 and NAC TF40 in Fig. 2D). Expression of GLK1, a transcription factor required specifically for M-cell differentiation in maize (Wang et al., 2013a) was M-enriched in our data as in prior RNA-seq studies (Fig. 5C).

MicroRNAs regulate mRNA stability and translation. To address whether cell-type specific microRNA expression accounts for the differential expression of any genes in BS and M cells, we sequenced small RNAs in the same RNA preparations used for RNA-seq. DESeq2 returned six differentially expressed microRNAs [|log_2_ (BS/M)| >1 and FDR < 0.05] (Supplemental Data S11), representing two microRNA families: MIR160 and MIR166 (Supplemental Fig. S8). Five members of the MIR160 gene family encode identical mature microRNAs, and these are strongly enriched in our BS fraction (Fig. 6A-B). One member of the MIR166 family (zma-MIR166a) (Supplemental Fig. S8) is enriched in our M fraction (Fig. 6A-B).

**Figure 6.**
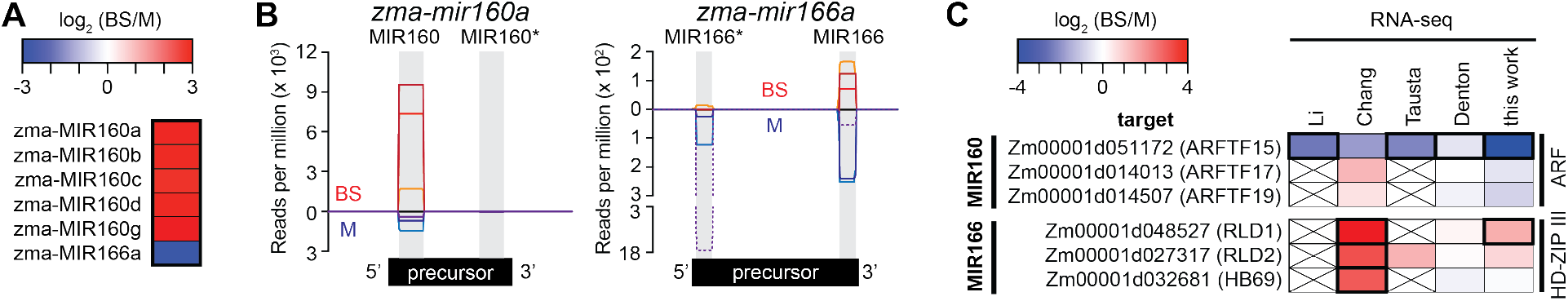
Differential expression of microRNAs and their transcription factor targets in BS and M fractions. (A) Differential expression of microRNAs from sRNA-seq analysis of BS and M fractions. Six genes identified by DESeq2 are shown [|log_2_ (BS/M)| >1 and FDR < 0.05]. The mature zma-MIR160 microRNAs have identical sequences, so sRNA-seq cannot distinguish them. (B) Normalized read coverage of genes encoding zma-MIR160a and zma-MIR166a. Coverage was normalized by million reads mapped to all microRNA precursors. Single sequence reads spanned the full length of each microRNA, resulting in uniform coverage across their length. Each line represents one replicate, with reads from the BS and M fractions shown above and below the horizontal line, respectively. Each microRNA precursor (diagrammed at bottom) folds into a hairpin via the pairing of complementary regions (gray). One strand in the duplex becomes mature microRNA and the other is typically degraded. An asterisk marks the degraded strand identified by miRBase (http://www.mirbase.org/). (C) Cell-type specific expression of mRNA targets of differentially expressed microRNAs. The mRNA targets of MIR166 and MIR160 were recovered from the DPMIND database (Fei et al., 2018). The original log_2_ (BS/M) values reported in prior RNA-seq datasets are summarized. Genes reported as differentially expressed are bordered in black. Crossed boxes indicate absence of data. ARFTF, auxin response transcription factor; HD-ZIP III, Class III homeodomain leucine zipper; RLD1/2, Rolled leaf1/2; HB69, Homeobox-transcription factor 69.

We then used the DPMIND database (Fei et al., 2018) to identify mRNA targets of these microRNAs. Zma-MIR160, which is enriched in our BS fraction, targets mRNAs encoding three auxin responsive factor (ARF) transcription factors. RNA-seq data show that expression of one of these, ARFTF15, is enriched in the M fraction (Fig. 6C), supporting a role for zma-MIR160 in directing ARFTF15 mRNA decay in BS cells. The M-enriched microRNA zma-MIR166 targets mRNAs encoding three HD-ZIP III transcription factors (Juarez et al., 2004; Fei et al., 2018). RNA-seq data reported by Chang et al (2012) indicate BS enrichment for all three of these mRNA targets, and our RNA-seq data support this for one of them (RLD1) (Fig. 6C). Zma-MIR166 is known to control spatial expression of RLD1 in maize leaf primordia (Juarez et al., 2004). Our data suggest further that zma-MIR166 regulates RLD1 in differentiated leaf tissue.

## Discussion

Numerous transcriptome studies have sought to identify genes whose differential expression is relevant to programming C_4_ traits (Wang et al., 2016; Schlüter and Weber, 2020). However, the contribution of translational regulation has been largely unexplored (Schlüter and Weber, 2020). Our study addressed this gap through analysis of BS and M translatomes in maize. Our results show that translational regulation contributes to the differential expression of genes in maize BS and M cells and provide clues about underlying mechanisms. Our data also add to the set of genes known to be differentially expressed in the two cell types, including new regulatory genes of potential relevance to the establishment or maintenance of C_4_ traits.

A conservative interpretation of our data identified 182 genes for which differential translation makes a strong contribution (>3-fold difference in TE) to their differential expression in BS and M cells (Supplemental Data S2). These include several genes encoding transcription factors and numerous genes encoding proteins involved in C_4_ photosynthesis, including one (the gene encoding RPI) that had previously been suggested to be under translational control (John et al., 2014) (Fig. 2D). Our previous study of chloroplast translatomes in BS and M fractions showed that controls at the level of translation and mRNA abundance often synergize to produce robust differential expression (Chotewutmontri and Barkan, 2016). Results here indicate that a similar theme holds true for many nuclear genes. The relative contributions of controls at the level of RNA abundance and translation vary and, in some cases, translational regulation plays the major role (Fig. 2B).

Ribo-seq can provide a more accurate view of differential expression than RNA-seq because it accounts for differences in both mRNA abundance and translational efficiency. In fact, our ribo-seq data provides evidence for the differential expression of many more genes than had been detected in prior RNA-seq studies (Fig. 5 and Supplemental Data S6 and S7). Of particular interest are new candidates for cell-type specific regulatory molecules. For example, we detected 44 new transcription factor candidates (Supplemental Data S10) including one (IAA10) that had been implicated in C4 differentiation on the basis of a cross-species selection scan (Huang et al., 2017), and two (NAC TF40 and THX28) that are expressed preferentially in the BS fraction due primarily to differential translation. Furthermore, we present evidence that the differential expression of two microRNAs underlies the cell-type enriched expression of several transcription factors: zma-MIR166a and zma-MIR160 are preferentially expressed in M and BS fractions, respectively, correlating with expression of their mRNA targets encoding transcription factors RLD1 and ARF proteins, respectively, in the opposite cell type (Fig. 6). Interestingly, a gain-of-function *Rld1* mutant that is no longer controlled by zma-MIR166 exhibited normal vasculature in leaf primordia (Juarez et al., 2004) but failed to increase vascular bundle density and lacked photosynthetic BS and M cells in mature leaf (Nelson et al., 2002). These results together with our findings suggest that zma-MIR166-mediated inhibition of RLD1 expression in mature M cells is important to maintain high vein density and differentiated BS/M cells, both of which are important C_4_ traits (Ermakova et al., 2020).

Our results also provide clues about the molecular basis for the differential translation events reported here. Our microRNA-seq data did not detect differentially expressed microRNAs that target translationally regulated mRNAs, suggesting that microRNAs do not contribute to translational control in this context. Our data hint, however, that differential use of AUG *versus* non-AUG initiation codons in uORFs is relevant to the cell-type dependent translation of several genes (Fig. 4C) and that non-AUG initiations are favored in BS cells. It is intriguing in this regard that specific eIF1 and eIF5A paralogs, two general translation factors that function in start codon selection (Nanda et al., 2009; Manjunath et al., 2019), exhibit BS-enriched expression (Fig. 4D). A method called Translation Initiation Site Profiling offers more robust detection of functional start codons than does ribo-seq, and revealed a shift to use of non-AUG start codons during yeast meiosis (Eisenberg et al., 2020). Analysis of the BS and M translatomes with this assay could address whether translational reprogramming of this type affects BS and M proteomes in biologically meaningful ways. In addition, sequence motifs that are enriched in UTRs of translationally regulated mRNAs (Fig. 3B) are candidate binding sites for sequence-specific RNA binding proteins. The motif UGUGCCG enriched in 3’-UTRs of mRNAs that exhibit preferential translation in BS cells (Fig. 3B) resembles the consensus binding site of PUF proteins, which regulate the translation of specific mRNAs in animals and fungi. Furthermore, the expression of one PUF-encoding gene is strongly enriched in the BS fraction (Fig. 3D), implicating this gene in the post-transcriptional control of gene expression in BS cells. PUF proteins typically repress translation, but examples of activation have been reported (Goldstrohm et al., 2018; Wang et al., 2018). Functional analysis of the candidate *cis*-elements and *trans*-factors to emerge from this study is an attractive direction for future investigation.

## Materials and Methods

### Preparation of BS and M fractions

We prepared BS and M fractions by using a minor modification of the rapid mechanical procedure described previously (Chotewutmontri and Barkan, 2016). In brief, *Zea mays* (inbred line B73) was grown for 13 days under cycles of 12 h light (300 μmol m^-2^ s^-1^) at 31°C and 12 h dark at 22°C. The apical one-third of the second and third leaves to emerge were harvested 2 h into the light cycle. The M fraction was taken as the supernatant following gentle grinding of fresh tissue in a mortar and pestle as described before (Chotewutmontri and Barkan, 2016). However, differing from our previous procedure which used the residual material from the M preparation as the source of the BS fraction, we prepared the BS fraction from separate tissue aliquots as the tissue remaining after several rounds of gentle grinding and rinsing to remove released cells. This fractionation procedure took roughly 3 min for M fractions and 10 min for BS fractions, at which point samples were flash frozen in liquid nitrogen.

### Ribo-seq, RNA-seq and sRNA-seq libraries

Ribosome footprints and total RNA were prepared from the same lysates in three biological replicates as described previously (Chotewutmontri and Barkan, 2016). Ribosome protected fragments from ∼20 to ∼40 nucleotides were selected for ribo-seq library preparation, and libraries were prepared with the NEXTflex Small RNA Sequencing Kit v2 (Bioo Scientific) in combination with rRNA depletion with customized biotinylated antisense oligonucleotides (Chotewutmontri et al., 2018). RNA-seq libraries were prepared from total RNA after rRNA depletion with the Ribo-Zero rRNA Removal Kit (Plant Leaf) (Illumina) using the NEXTflex Rapid Directional qRNA-Seq Kit (Bioo Scientific). To catalog microRNAs, the same RNA samples were analyzed by sRNA-seq with the same kit used to sequence ribosome footprints. The ribo-seq, RNA-seq and microRNA libraries were sequenced using a HiSeq 4000 (Illumina) in single-read mode with read lengths of 100 nucleotides.

### Sequence read processing

Ribo-seq and RNA-seq data were processed as described previously (Chotewutmontri and Barkan, 2020) with minor modifications. In brief, adapter sequences were trimmed using cutadapt (Martin, 2011). For ribo-seq, only trimmed reads between 18 and 40 nucleotides were retained for alignments. Reads were aligned sequentially to the maize chloroplast genome (GenBank accession X86563), the maize mitochondrial genome (B73 RefGen_v4 assembly release 38), and the maize nuclear genome (B73 RefGen_v4 assembly release 38) using STAR version 2.5.3a (Dobin et al., 2013). Read splicing was restricted to annotated splice junctions. Read counting was performed using featureCounts from Subread package version 1.6.0 (Liao et al., 2014), counting only those reads that mapped to a unique site. Ribo-seq read counts for nuclear and organellar genes excluded the first 25 and 10 nucleotides of each ORF, respectively, to avoid counting ribosome pileups at start codons. RNA-seq read counts used full length ORFs. Read counts are provided in Supplemental Data S12.

Ribosome footprint length distribution, three-nucleotide periodicity, and metagene analysis were performed using in-house scripts. Comparison among replicates was evaluated with Pearson correlation and hierarchical clustering using log_10_ RPKM values. Transcript coverage and 3’ bias were analyzed using Picard Tool version 2.6.0 (http://broadinstitute.github.io/picard/). Samtools (Li et al., 2009) was used to extract gene-specific read coverages. P-site assignment of ribo-seq reads was inferred from the ribosome footprint size-dependent placements of footprint 5’ ends to the P-site location, which were observed from metagene analysis of reads mapping to start codons in the data discussed here.

MicroRNA data were trimmed with cutadapt (Martin, 2011). Trimmed reads between 15 and 45 nucleotides were retained and aligned to the B73 RefGen_v4 assembly using STAR version 2.5.3a (Dobin et al., 2013). For multi-mapped reads, STAR parameters were set to distribute each multi-mapped read randomly to one of the mapped locations (--outMultimapperOrder Random --outSAMmultNmax 1). Read counting was performed using featureCounts (Liao et al., 2014) with maize microRNA annotation release 22 (mirbase.org), where counting included multi-mapped reads. The read counts are provided in Supplemental Data S11.

### Differential expression analysis

Differential expression analysis based on RNA-seq, ribo-seq or microRNA data was performed using DESeq2 (Love et al., 2014). Outputs are reported in Supplemental Data S5, S11 and S13. Genes were reported as differentially expressed if they showed an average of at least 100 reads in at least one of the tissue types, a |log_2_ BS/M| > 1 and FDR < 0.001 for RNA-seq and ribo-seq or FDR < 0.05 for microRNA data.

For analysis of differential TE, we used XTAIL (Xiao et al., 2016). Only genes with an average of at least 100 RNA-seq reads in both tissue types and an average of at least 100 ribo-seq reads in at least one of the tissue types were reported. Genes were reported as differentially translated if they showed |log_2_ (BS/M) TE| > 1.585 with FDR < 0.001. The XTAIL output is provided in Supplemental Data S1.

### UTR sequence motif analysis

UTRs from genes that are differentially translated (107 genes in BS set and 75 genes in M set; |log_2_ (BS/M) TE| > 1.585 excluding the buffering genes) were compared with one another and with a set of non-differentially translated genes (Control set; 5280 genes; |log_2_ (BS/M) TE| ≤ 0.585). The UTR sequences were downloaded from the BioMart of Ensembl Plants release 50 (https://plants.ensembl.org/). Missing UTR sequences for genes in the BS and M sets were manually extracted from GenBank. Analyses used the longest annotated UTR sequences for each gene. Statistical testing was performed using Prism 8 (GraphPad). Motif discovery was performed using STREME (Bailey, 2020) with a motif width of 4 to 25 nucleotides. Motifs with p-values < 0.05 were retained. To remove known DNA motifs and identify known RNA motifs, the motifs were searched against three DNA and RNA motif datasets (JASPAR CORE, CISBP-RNA Zea mays and Arabidopsis DAP motifs) using TOMTOM (Gupta et al., 2007) based on Euclidean distance. No known RNA motifs were detected. DNA motifs that produced any match with a p-value below 0.07 were removed. Motif enrichment was performed using AME (McLeay and Bailey, 2010). FIMO (Grant et al., 2011) was used to find motif positions in UTR sequences and only the top score FIMO sites corresponding to the number of sites found by AME were used. The motif analysis data are provided in Supplemental Data S3.

### uORF analysis

Genes with |log_2_ (BS/M) TE| > 1 and FDR < 0.001 excluding the buffering genes (n=335) were screened manually for ribo-seq reads in the 5’UTR. P-site assignments and identification of uORFs beginning with AUG, CUG or ACG were determined for those genes with substantial ribo-seq reads in the UTR. Genes that show P-site coverage in predicted uORFs are reported in Supplemental Data S4. uORF read count was performed using featureCounts (Liao et al., 2014), counting only those reads that mapped to a unique site and excluding the 25-nucleotide region upstream of the main ORF to avoid counting ribosome pileups at start codons. The TE of uORFs was calculated from the normalized ribo-seq abundance (number of ribo-seq reads mapped to the uORF per uORF kilobase per million reads mapped to nuclear coding sequences) divided by the normalized mRNA abundance (mORF RNA-seq RPKM).

### Functional enrichment analysis

The enrichment analysis was performed with MapMan (Thimm et al., 2004) as described (Zones et al., 2015), using maize X4.2 functional assignments and Mapman terms Levels 1 to 3. In brief, a set of genes showing an average ribo-seq RPKM value > 1 in at least one tissue type (18,980 genes) was used to calculate background distribution of each MapMan term among the expressed gene set. To evaluate a pair of BS and M data sets of sizes *m* and *n* genes, a total of 10,000 permutations by random sampling without replacement for pair sets of *m* and *n* genes from the expressed gene pool were performed. The term occurrences in these 10,000 permutated data set pairs produced the mean and standard deviation for each term occurrence in the background. The observed term occurrence in each data set together with the mean and standard deviation of the term occurrence in the background from the corresponding permutation set were used to calculate z-score and p-value. The adjusted p-values or FDR values were calculated using the Benjamini-Hochberg method in R as described previously (Zones et al., 2015). The output is provided in Supplemental Data S8.

### Reanalysis of external data

Previously published RNA-seq analyses of maize BS and M fractions used non-current maize genome assemblies and gene models. For comparison to our data, we re-analyzed the +9 cm data (also called section 14) from Tausta et al. (2014) and the slice 1 data from Denton et al. (2017) using our pipeline. To compare our ribo-seq data to proteome data from Friso et al. (2010), those data were matched to B73 v4 gene identifiers using the gene ID history (gramene.org). For comparison to our differential translatomes in Figures 3E, 5, 6 and Supplemental Figure S6 and S7, the originally reported BS/M values were compiled from the BS and M data from Li et. al. (2010), Chang et al. (2012), the section 14 data from Tausta et al. (2014), and the slice 1 data from Denton et al. (2017). The values reported with older gene IDs were matched using the gene ID history (gramene.org).

### Antibodies

Antibodies to NdhH and PPDK were generously provided by Tsuyoshi Endo (Kyoto University) and Kazuko Aoyagi (University of California, Berkeley), respectively. Antibodies to PEPC, malic enzyme, and RbcL were gifts of William Taylor (University of California, Berkeley). The antibody to D2 was obtained from Agrisera.

### Accession Numbers

The ribo-seq and RNA-seq data were deposited at the NCBI Sequence Read Archive (SRA) with accession number PRJNA667075. RNA-seq data from Tausta et al. (2014) and Denton et al. (2017) were downloaded from SRA (accession numbers SRP035577 and SRP052802, respectively). Alignments of reads to the maize chloroplast genome used Genbank accession X86563. B73 RefGen_v4 assembly with annotation release 38 (gramene.org) was used for alignments to the nuclear and mitochondrial genomes.

## Supplemental Material

Supplemental Figure S1. Characteristics of ribo-seq and RNA-seq data from BS and M fractions.

Supplemental Figure S2. Comparison of RNA-seq data characteristics with those from prior studies.

Supplemental Figure S3. Comparison of RNA-seq, ribo-seq, and proteome data collected for maize BS and M leaf fractions.

Supplemental Figure S4. Comparison of our RNA-seq and prior RNA-seq data for genes exhibiting preferential translation in M or BS fractions.

Supplemental Figure S5. Normalized read coverage of translationally regulated genes. Supplemental

Figure S6. Differential expression of eIF1, eIF5 and eIF5A genes.

Supplemental Figure S7. Differential expression of biogenesis factors for PSII, Rubisco, and the NDH complex.

Supplemental Figure S8. Alignment of microRNA sequences of zma-MIR160 and zma-MIR166 families.

Supplemental Data S1. XTAIL output and related data.

Supplemental Data S2. Genes reported by XTAIL as having a greater than three-fold difference in TE between BS and M fractions.

Supplemental Data S3. Sequence motif enrichment data.

Supplemental Data S4. Data on uORF translation for differentially translated genes Supplemental Data

S5. DESeq2 output of ribo-seq data.

Supplemental Data S6. BS-enriched genes in ribo-seq data. Supplemental

Data S7. M-enriched genes in ribo-seq data.

Supplemental Data S8. Enrichment of MapMan terms in gene sets that are differentially expressed in BS versus M fractions based on ribo-seq data.

Supplemental Data S9. Differential expression of nucleus-encoded proteins involved in expression and assembly of Rubisco, NDH-like and PSII complexes.

Supplemental Data S10. Data for transcription factor genes that are differentially expressed based on ribo-seq data.

Supplemental Data S11. MicroRNA data.

Supplemental Data S12. Ribo-seq and RNA-seq read counts, including re-analyzed data from Denton S1 and Tausta Sec 14 samples.

Supplemental Data S13. DESeq2 outputs of RNA-seq data, including re-analyzed data from Denton S1 and Tausta Sec 14 samples.

## Acknowledgments

We wish to thank Rosalind Williams-Carrier for assistance with sample preparation and for comments on the manuscript, Susan Belcher for comments on the manuscript, the University of Oregon Genomics and Cell Characterization Core Facility for Illumina sequencing, and the University of Oregon Research Advanced Computing Services for access to high performance computer, Talapas. This work was supported by grant IOS-1339130 to A.B. from the US National Science Foundation.

